# From sensation to action: distributed mesoscale dynamics in the primate brain

**DOI:** 10.64898/2026.10.04.756523

**Authors:** Sunhang Shi, Meixuan Chen, Zhiyan Quan, Chenxi Zhai, Meizhen Qian, Hisashi Tanigawa, Ke Jia, Xiaotong Zhang, Gang Pan, Anna Wang Roe

## Abstract

Perceptual decisions engage distributed brain networks, yet how decision-related signals are organized across and within brain areas remains unclear. Mesoscale functional domains are established in visual cortex, but whether spatially distinct domains organize perceptual decisions across the brain is unknown. Here we used 7-T functional MRI at 1-mm isotropic resolution in two awake macaques performing a contrast-discrimination task to map brain-wide activity during sensation, decision formation and saccadic report. Voxel timecourses separated into three groups with distinct temporal profiles, associated with sensory, decision-related and action-related activity. Their spatial distributions formed a broad progression from visual cortex to parietal, prefrontal and motor regions, while revealing distinct response domains within these regions. Choice-related signals were distributed across sensory, associative, and motor systems before movement, including in visual cortex responses to physically identical stimuli that differed according to the animal’s eventual choice. Moreover, within the intraparietal sulcus and other regions, mesoscale domains preferentially responsive to near-threshold conditions carried choice information under uncertainty, whereas domains responsive to high-contrast conditions carried choice information when evidence was strong. These findings reveal that perceptual decisions are organized across the brain through spatially differentiated mesoscale networks whose activity varies with task phase and sensory certainty. This spatial organization provides a framework for investigating how distributed neural signals support flexible choice.

## Introduction

Historically, the study of brain function has emerged from modality-specific approaches, leading to studies categorized as, for example, circuits underlying sensory, motor, cognitive, or limbic behavior.^1^However, our behaviors are wholistic, whereby each behavior exhibits components of each of these modalities. As is often true in science, availability of new technology leads to shifts in how we address fundamental questions and to shifts in hypotheses that can be practically addressed. More specifically, recent years have seen the rapid proliferation of methods enabling the recording of thousands of neurons simultaneously and, furthermore, across many regions of the brain^2,3^. This has led to a flurry of exciting publications emphasizing the distributed brainwide nature of the neuronal code^4–10^, leading to re-evaluation of well-studied behaviors such as decision making. While these studies have offered recognition of neuronal contributions from many areas of the brain, there remains a lack of organizational understanding regarding how these many areas contribute to specific behaviors.

Here, we approach distributed systems from a functional organizational perspective. Our working hypothesis is that each cortical area represents a distinct axis of behavior and uses its intracortical circuitry for computing values within its parameter set. We explore the hypothesis that different values may be represented in distinct mesoscale spatial locations, much as it is in visual cortical areas. To address this hypothesis, we have turned to ultrahigh field MRI technology. Functional MRI offers a complementary approach to electrophysiology. Its key strength is mapping brainwide activity, which coupled with recent advances in ultrahigh field technology, attains millimeter-scale resolution^11,12^. In nonhuman primates, we and others have pioneered this approach by developing methodologies for studying brain function in monkeys [cite others], and, in particular, in macaque monkeys performing behavioral tasks in 7T MRI (RF coils^13,14^, modifications of analysis pipelines to remove steps that introduce blur^15,16^, novel animal behavioral training methods^17^, MR-compatible brains stimulation methods^18–20)^. These have demonstrated that mesoscale precision is needed to probe how the brain represents information, not only in sensory areas but across the brain^21^. These studies show that higher resolution is not just refinement of current understanding; it provides qualitatively distinct understanding of functional specialization within areas, leading to maps of distinct variables represented within single areas. If task-relevant variables occupy distinct mesoscale domains, they may be accessible through relatively simple, spatially selective readout. This possibility provides a computational motivation for studying the spatial organization of distributed brain activity.

To test our hypothesis regarding an organized functional view of behavior, our testbed is the classic decision-making task, comprising sensory, decision making, and response phases, performed by trained monkeys. Previous studies have identified widespread correlates of perceptual decisions and distinguished response profiles associated with sensory processing, evidence accumulation and behavioral reports^22–27^. Electrophysiological studies have provided signals for sensory evidence in early visual cortex and the middle temporal area (MT), evidence-related ramping activity in parietal (notably lateral intraparietal area (LIP)^28–32)^ and frontal circuits, and signals associated with motor commitment in the frontal eye fields(FEF)^33–35^ and superior colliculus(SC)^36–38^. These findings support models that link sensory encoding, evidence accumulation, and action.

Here, we predict that ultrahigh field fMRI will reveal, for each phase of the task, activations in the previously identified areas (such as visual cortex and intraparietal sulcus). We also predict mesoscale specification with each area, reflecting spatially distinct response values of variables such as sensory evidence and choice representation. Experimentally, we combined 7T fMRI with a two-alternative contrast discrimination task in two awake macaques to map distributed mesoscale activity. By identifying voxels related to stimulus viewing, decision making, and saccadic report, we found shifts in global (within parietal, prefrontal and motor regions) BOLD-based networks across these phases. Locally, within areas, mesoscale spatial shifts related to the variable of sensory certainty were also observed. Together, these findings reveal a mesoscale functional organization linking sensation to action within and across areas of the primate brain, providing a spatial framework for understanding how biological intelligence flexibly translate uncertain sensory information into action and for developing efficient, spatially informed readout strategies for brain– computer interfaces.

## Results

Here, we present behavioral results from two monkeys performing a two-alternative force choice contrast discrimination task in a 7T MRI (Fig. 1). We describe the behavior, and how the mesoscale BOLD activations during the sensory, decision making, and action phases of the task are defined (Fig. 2). This leads to a brain-wide view of representation of choice (Fig. 3). We then provide an analysis of the networks associated with different degrees of task certainty (Fig. 4).

**Fig. 1.**
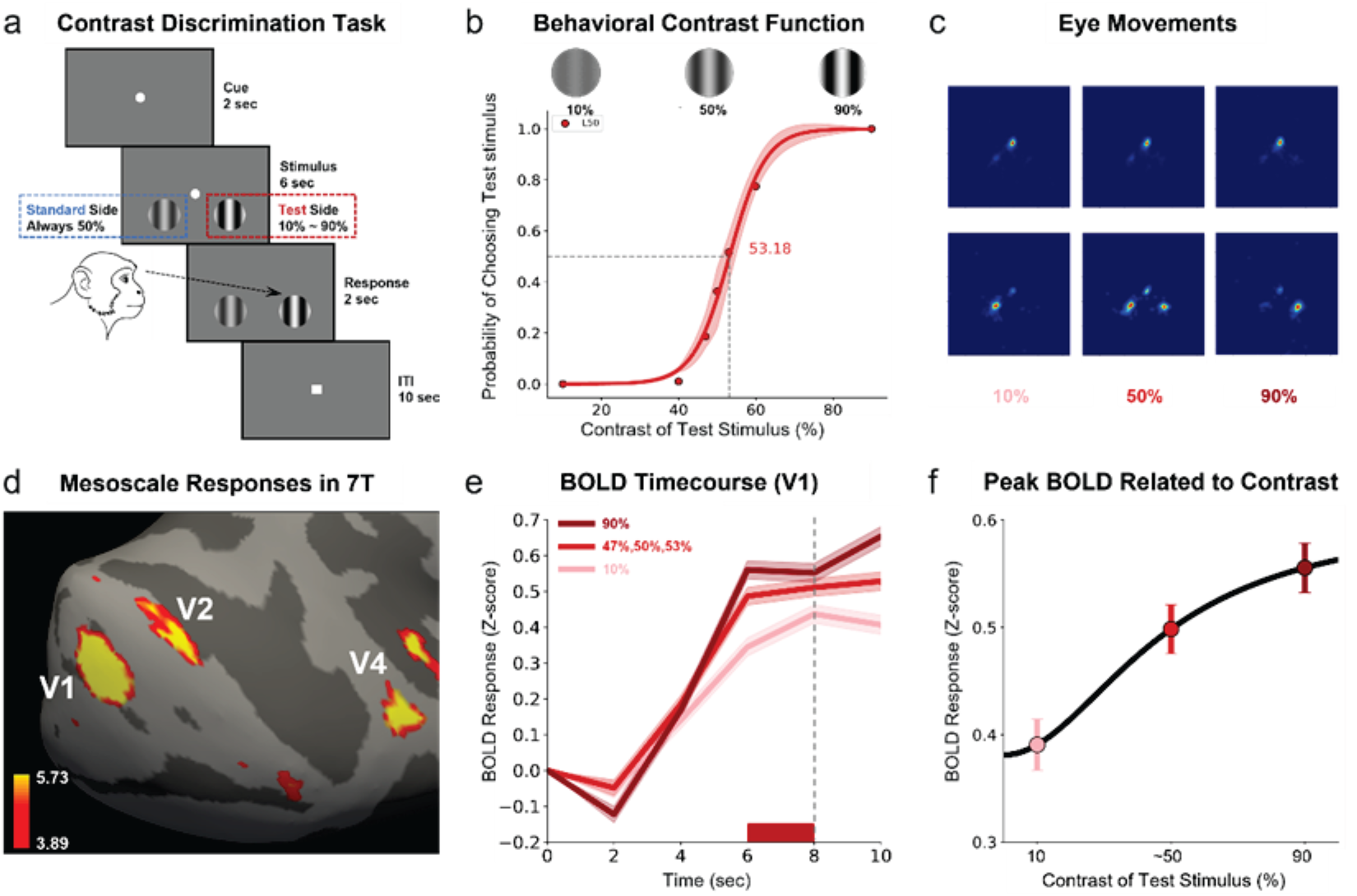
Contrast discrimination paradigm and mesoscale fMRI mapping. **a,** Each trial began with central fixation (2 s), followed by bilateral grating stimuli (6 s) where *the standard side* (blue dashed box) was fixed at 50% contrast and *the test side* (red dashed box) contrast varied across trials (10%, 40%, 47%, 50%, 53%, 60%, 90%). At fixation dot offset, subjects responded by making an eye saccade to the higher contrast stimulus within a 2 s response window. Trials were separated by a 10 s inter-trial interval (ITI). **b,** Representative psychometric function showing the probability of choosing the test stimulus (monkey M; n = 5 sessions, 466 trials, the left side was fixed as the standard stimulus). At low test contrast (e.g., 10%), subjects predominantly chose the standard side, at high test contrast (e.g., 90%), they reliably chose the test side. When test contrast was close to 50%, choices approached chance level (∼50%), reflecting increased perceptual ambiguity. The point of subjective equality (PSE) was estimated at 53.18% for monkey M and 51.86% for monkey J, under conditions where the left side was fixed as the standard stimulus. Shadowed error bars represent the standard error of the mean (SEM) across all trials for each of the seven test contrast levels (10%-90%). **c,** Eye position heatmaps for three representative contrast levels (10%, 50%, 90%) showing tight fixation during stimulus viewing (above) and contrast-dependent saccade endpoints during the response phase (below). At 50% contrast, the bimodal distribution reflects higher task difficulty. These results rule out eye movement confounds during early sensory encoding. **d,** Signal difference in mesoscale functional organization in early visual cortex mapped in 7T fMRI during a fixation task. The color bar is indexed by corresponding Z scores, where 3.89 and 5.73 represent *FDR-corrected p* < 10⁻^4^ and *p* < 10⁻^8^, respectively. **e–f,** Contrast-dependent BOLD responses in V1 measured via mesoscale fMRI. **e**, Time course of BOLD signals from V1 across test contrast conditions. Responses increased monotonically with contrast, from pink (10%) through red (average of 47%, 50%, and 53%) to dark red (90%). Shaded areas denote SEM across trials. The gray dashed vertical line marks the onset of saccade initiation. **f**, Peak BOLD responses (averaged over 6–8 s, red horizontal bar in Fig. 3e) plotted as a function of test stimulus contrast. Data were well-fit by a Naka–Rushton function, capturing the characteristic compressive nonlinearity of cortical contrast gain. Error bars indicate the standard error of the mean (SEM) across trials. Valid trial counts at each contrast level were as follows: 65 trials at 10%, 67 at 90%, and 207 combined across 47%, 50%, and 53% contrasts. All data were from monkey M under the condition where the left side was fixed as the standard stimulus.

**Fig. 2.**
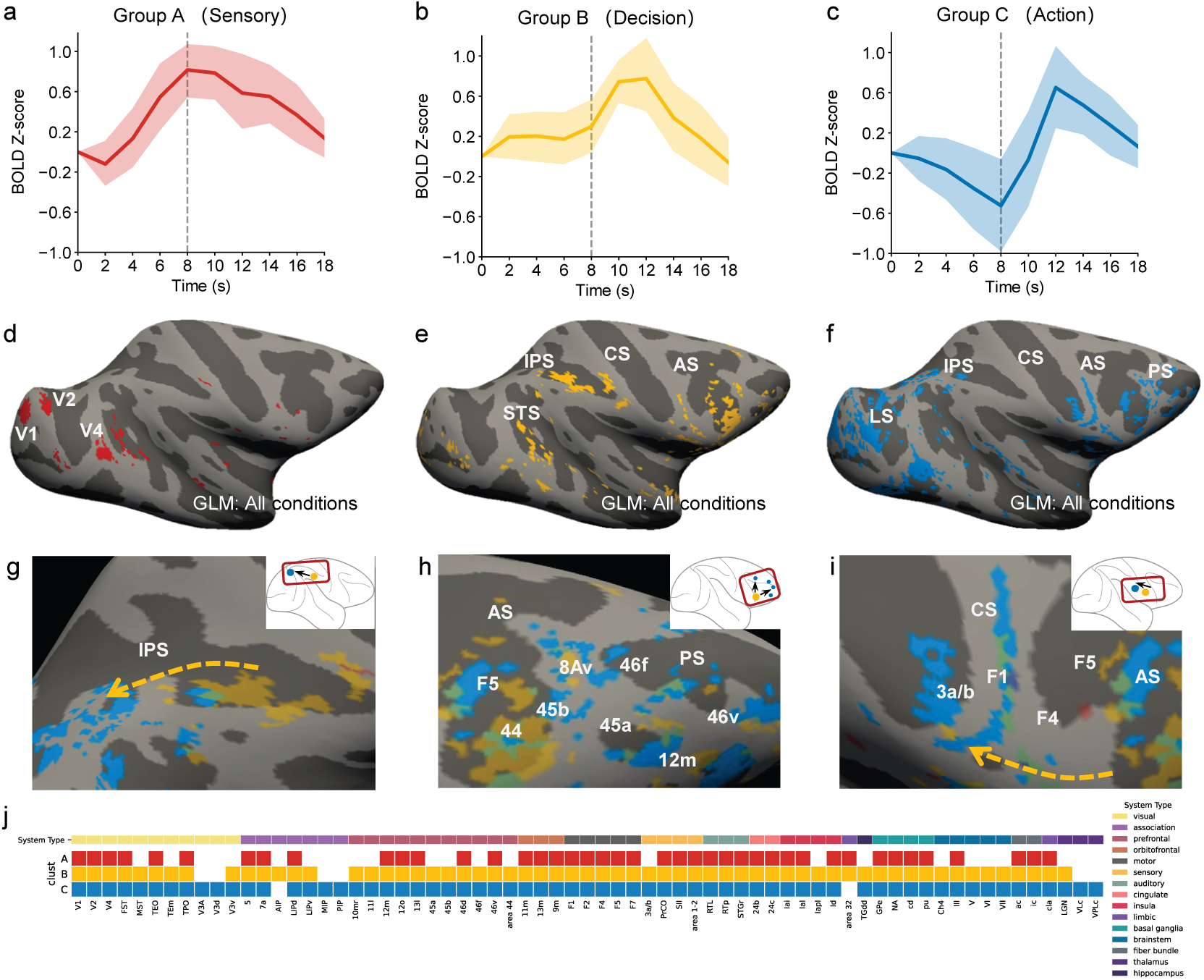
Spatial transition of three functional brain-wide networks identified from timecourse. **a–c,** Voxel-averaged BOLD time courses for the three groups. Significant voxels were identified using FDR*-corrected p* < 0.01 for the fixation regressor and FDR*-corrected p* < 10⁻^6^ for the response regressor. For each significant voxel, BOLD time courses were then averaged across all conditions and trials. These voxel-level time courses were subsequently averaged within each group (group A, red, N = 455; group B, yellow, N = 1782; group C, blue, N = 3617). Shaded regions indicate the interquartile range (IQR; 25th–75th percentiles) across voxels within each group. **d–f,** Spatial Distributions of three groups with distinct signal patterns on the cortical surface. **d,** Voxels categorized as group A (red; fixation regressor with steep slope) localized primarily to early visual areas. **e,** Voxels categorized as group B (yellow; fixation regressor with gradual ramping slope) were distributed across intraparietal sulcus (IPS), superior temporal sulcus (STS), and lateral prefrontal cortex. **f,** Voxels categorized as group C (blue; significantly modulated by the response regressor) were concentrated in premotor and frontal regions. All GLM estimates were derived from models incorporating all contrast conditions (10%, 40%, 47%, 50%, 53%, 60%, 90%). **g-i**, Magnified view of the IPS(g), PFC(h) and motor area(i) showing local transitions from group B to group C, respectively. Yellow patch: group B; blue patch: group C; yellow arrow: indicating the network shift. Inset (top right): schematic localizer of the whole-brain surface; red box corresponds to the magnified region, yellow-to-blue arrow indicates topographic shift. **j**, Area-by-category matrix. Each row represents a group with distinct signal pattern— group A (red), group B (yellow), or group C (blue)—as defined by GLM regressor identity and fixation-period slope. Each column corresponds to an area defined in the D99 macaque brain atlas. Areas are grouped by system affiliation (IPS, intraparietal sulcus; PFC, prefrontal cortex; color bar, top). For visualization, only areas containing more than 10 voxels were included.

**Fig. 3.**
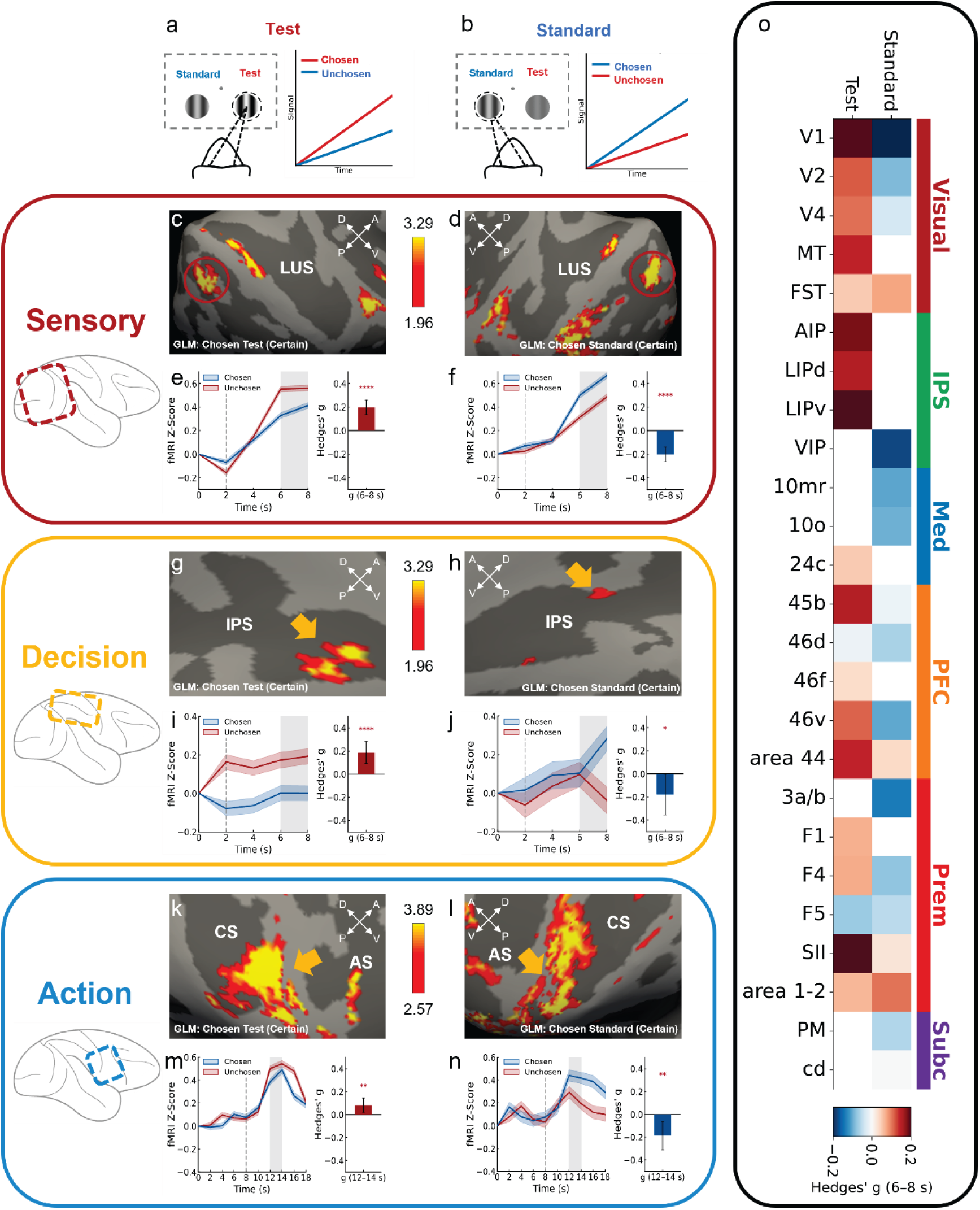
Choice representation encoding under high-certainty conditions. **a–b,** Schematic illustration of the chosen–unchosen comparison used to assess choice representation under high-certainty conditions. **a,** High-certainty trials in which the monkey chose the high-contrast test stimulus (90%). When the test stimulus was chosen (red), its contralateral cortical response was higher than when the test stimulus was unchosen. **b,** High-certainty trials in which fixed 50% standard stimulus was chosen. When the standard stimulus was chosen (blue), the contralateral hemisphere showed stronger responses compared with unchosen standard trials. **c–f,** Example from the sensory group (group A), using V1 as a representative case, showing clear chosen–unchosen modulation consistent with a choice representation signal. Representative data was from monkey M (90% contrast: n = 65 trials; 10% contrast: n = 63 trials; standard stimulus fixed at 50%). Schematic locators indicate the position of magnified area relative to the whole brain, with red boxes denoting the zoomed-in regions. **c,** Hemisphere contralateral to the test stimulus. Voxels classified as the sensory group are shown on inflated cortical surfaces. LUS, lunate sulcus. The color bar is indexed by corresponding Z scores, where 1.96 and 3.29 represent *FDR-corrected p* < 0.05 and *p* < 0.001, respectively. **d,** Hemisphere contralateral to the standard stimulus. **e–f.** Averaged BOLD responses from V1 voxels which were activated when test stimuli was chosen, shown separately for the hemisphere contralateral to the test stimulus (e) and to the standard stimulus (f). For each panel, the left plot shows mean time courses of BOLD responses, separated by whether the test stimulus (red) or the standard stimulus (blue) was chosen. The right plot shows standardized effect sizes for the 6–8 s fixation window, quantified by Hedges’ g (mean ± 95% bootstrap confidence interval), providing a bias-corrected measure of the chosen–unchosen difference that is comparable across ROIs with different voxel counts. In both contrast conditions, V1 responses in the contralateral hemisphere were significantly higher on chosen than on unchosen trials (two-sample Welch’s t-test on trial-wise 6– 8 s BOLD averages; significance denoted as: **** for *p* < 10⁻⁴, *** for *p* < 10⁻³, ** for *p* < 0.01, * for *p* < 0.05; no marker indicates non-significance). **g–j,** Demonstration of decision-related responses in IPS (decision group, group B). Voxels classified as the decision group (*FDR-corrected p* < 0.05) are shown on inflated cortical surfaces. Activations in the hemisphere contralateral to the test stimulus (g) or the standard stimulus (h), under high-certainty conditions. IPS, intraparietal sulcus. **i,** Example BOLD responses from IPS in the hemisphere contralateral to the test stimulus. Left, mean time courses separated by chosen-test (red) and chosen-standard (blue) trials. Right, corresponding effect sizes (Hedges’ g) for the 6–8 s fixation window. **j,** Hemisphere contralateral to the standard stimulus. In both regions, contralateral responses were stronger on chosen versus unchosen trials, indicating a reliable choice-related encoding in decision group. **k–n,** Demonstration of action-related responses in action group (group C). Response-period BOLD activations in motor cortex for hemispheres contralateral to the test stimulus (k) or the standard stimulus (l) under high-certainty conditions. CS, central sulcus; AS, arcuate sulcus. The color bar is indexed by corresponding Z scores, where 2.57 and 3.89 represent *FDR-corrected p* < 0.01 and *p* < 10⁻⁴, respectively. **m,** Example BOLD responses from action group in the hemisphere contralateral to the test stimulus. Left, mean time courses separated by trials in which the test stimulus was chosen (red) versus trials in which the standard stimulus was chosen (blue). Right, corresponding effect sizes (Hedges’ g) for the response-period window. In both cases, contralateral responses were higher when the corresponding stimulus was chosen than when it was unchosen, indicating a reliable choice-related encoding in action group. **n,** Response from hemisphere contralateral to the standard stimulus. **o,** Summary matrices of chosen–unchosen effect sizes (Hedges’ g, 6–8 s) across all ROIs prior to the response period, spanning both sensory and decision groups. Each row corresponds to one ROI, grouped and color-coded by functional system. Color intensity reflects the magnitude of the chosen–unchosen difference, quantified using Hedges’ g (6–8 s window). Only ROIs with at least three such voxels were included. System labels are abbreviated as follows: Med = Medial, Prem = Premotor, and Subc = Subcortical.

**Fig. 4.**
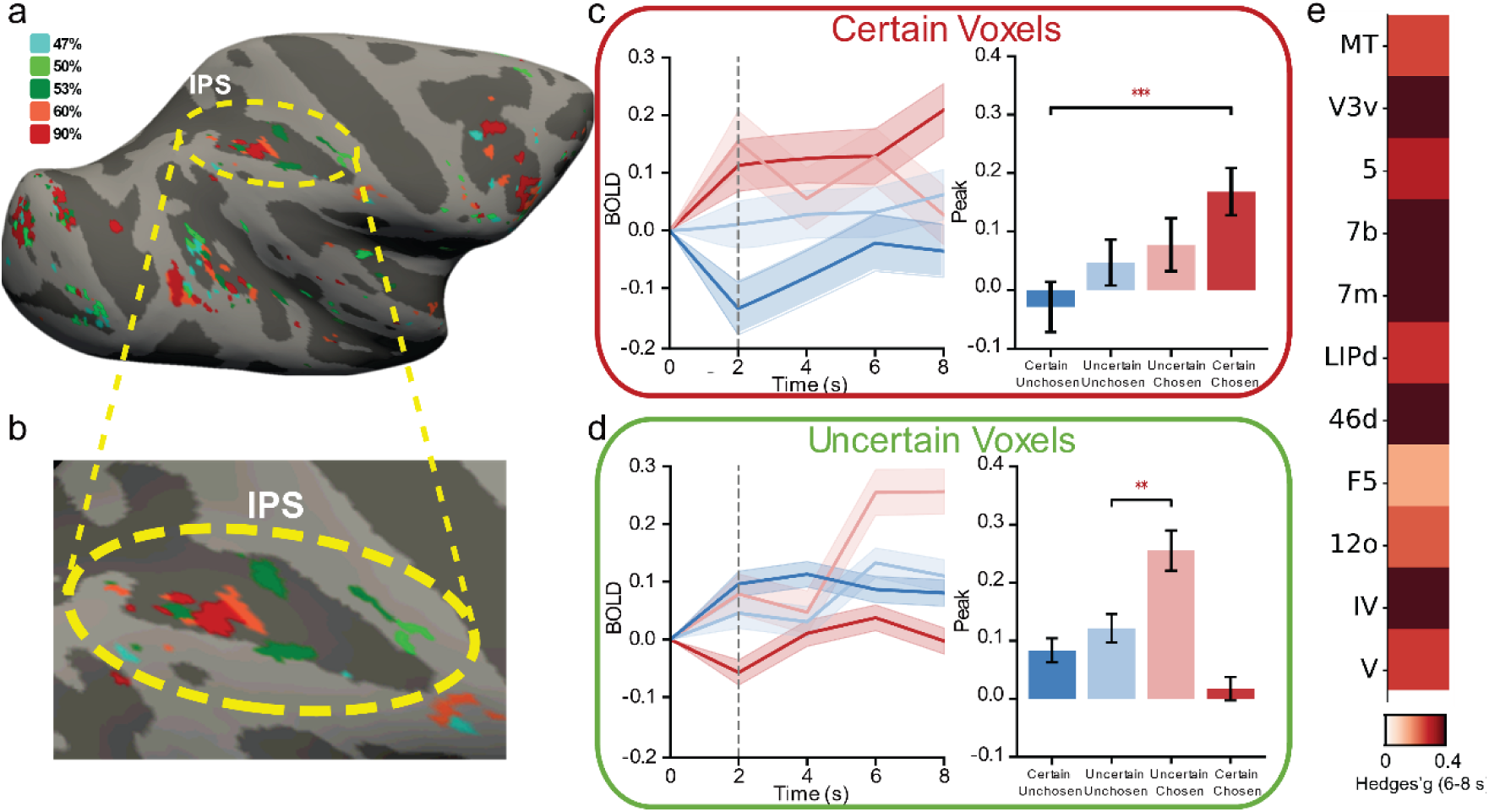
Uncertainty encoding by mesoscale specificity. **a**, Winner-take-all (WTA) map across the cortical surface. For each voxel, we estimated a condition-wise GLM and first thresholded each condition’s (*FDR-corrected p* < 0.05). Among the surviving voxels, the condition with the minimum q value was assigned as the voxel’s “winning” condition. Voxels were color-coded by the winning conditions: green for near-threshold contrasts (47%, 50%, 53%), defined as uncertain conditions where behavior approached chance performance; and red for large contrast differences (60%, 90%), defined as certain conditions where behavioral choices reliably favored the test stimulus. The resulting voxel-wise labels were projected onto the surface to obtain a brain-wide mesoscale WTA map. **b**, Example region in the intraparietal sulcus (IPS). For further analyses, we selected voxels with minimum q values at 90% contrast (deep red) to represent certain test-chosen voxels, and voxels with minimum q values at 47% or 53% (green) to represent uncertain voxels. **c**, Amplitude responses for IPS certain voxels. Bar plots show mean BOLD amplitude for trials in which the test stimulus was chosen (red) or not chosen (blue). Deep red and deep blue bars correspond to certain conditions (90% chosen test; 10% unchosen test), whereas pale red and pale blue bars correspond to uncertain conditions (47–53%). Certain voxels showed significantly higher responses for trials in which the test stimulus was chosen than not chosen during certain conditions, but showed no such difference during uncertain conditions. **d**, Amplitude responses for IPS uncertain voxels. For voxels identified as uncertain (47% and 53%) in WTA maps, the pattern reversed: certain trials (10%, 90%) showed no amplitude separation, whereas uncertain trials exhibited significantly stronger responses for trials in which the test stimulus was chosen than unchosen. Thus, uncertain voxels form a distinct network that remains informative under uncertain conditions, allowing choice encoding even when sensory evidence is ambiguous. **e**, Across cortical regions, we quantified whether WTA-defined uncertain voxels could predict choices under uncertain conditions. The resulting matrix summarizes the choice-encoding strength of uncertain voxels across cortical regions.

Although BOLD responses unfold over several seconds, the neural events that drive them occur at particular times and in distinct spatial locations. By combining high spatial resolution with whole brain coverage, mesoscale fMRI can therefore use spatially distributed activity to reveal the progression of neural processing across task phases, despite the slow temporal response of BOLD.

### Contrast-dependent responses in early visual cortex

We hypothesized that, within a standard 2-AFC task, different phases of the task engage distinct brain-wide mesoscale networks. During sensory phase of the task, we predicted that mesoscale BOLD responses in early visual cortex would exhibit contrast-dependent contrast response functions (CRFs) that reflect the strength of sensory evidence during perceptual decision-making. In particular, we expected signal amplitudes to increase with stimulus contrast, and sought to determine whether these signals also carry choice-related information under conditions of matched sensory input.

#### Behavioral paradigm

To examine the brain-wide representation of sensory, decision-related, and motor signals, we trained macaques to perform a two-alternative forced choice (2AFC) contrast discrimination task in 7T MRI (Fig. 1a). This well-established paradigm has been widely used in electrophysiological studies^2^ to dissociate the neural correlates of sensory encoding, decision formation, and action execution.

Each trial began with a 2-second central fixation cue, followed by 6 seconds (sufficient time for capturing the BOLD signal) of bilateral drifting grating presentation (at 4.2deg eccentricity) during which the monkey maintained fixation within a 2°-radius window centered on a 0.2°-radius fixation point (Fig. 1A). Upon cue offset, the animal indicates the side with the higher perceived contrast by making a saccade to the corresponding grating stimulus (within a 1°-radius stimulus). Correct responses were rewarded with juice. A trial was considered valid if the subject maintained central fixation during stimulus presentation and executed a saccade during the response period.

We collected a total of 1,761 valid trials across 18 sessions from two animals (monkey M: 875 trials; monkey J: 886 trials). In each session, the stimulus on one side (the *standard stimulus*) was fixed at 50% contrast (for each day presented in one hemifield; left or right hemifields counterbalanced across days), while *the test stimulus* appeared in the opposite hemifield with contrast levels pseudo-randomly selected from values between 10% and 90% (10%, 40%, 47%, 50%, 53%, 60%, 90%). In a single session, the probability that the higher-contrast stimulus appeared on the left side was equal to the probability that it appeared on the right side.

Behavioral performance on this task followed a classic sigmoidal psychometric function (Fig. 1b, monkey M; Fig. S1a, monkey J), with choice probability increasing as test contrast rose relative to the fixed 50% standard. At low test contrast (e.g., 10%), animals reliably chose the standard side, while high test contrast (e.g., 90%) drove choices consistently toward the test side. At intermediate test contrasts (∼50%), performance approached chance. This performance was consistent with greater certainty for high- and low-test contrasts and decreased decision certainty for intermediate values. The point of subjective equality (PSE) was estimated at 53.18% for monkey M and 51.86% for monkey J (Fig. S1), consistent with contrast-based discrimination and near-symmetric performance around the 50% standard. By monitoring the eye movements of the monkeys, we confirmed that animals maintained stable fixation during stimulus presentation and executed saccades in the response phase (Fig. 1c, monkey M). Eye position heatmaps and time traces showed minimal movement during the fixation period, ruling out eye movement confounds in early sensory responses. During the response epoch, saccades became more variable and often bimodal under low contrast differences, reflecting increased task difficulty. Similar results were obtained for the monkey J (Fig. S1). The classic behavioral contrast functions and the eye movements patterns illustrate that the monkey’s behavior reflects the expected choice task response and is stable and reliable.

#### fMRI procedures

A fundamental constraint of fMRI in perceptual decision-making research is the extended temporal rise of the BOLD signal, which precludes direct measurement of sub-second neural dynamics. Despite this slow rise, the hemodynamic signal following a sensory or cognitive event can signal the occurrence and the intensity of that event; this association has formed the basis of using functional imaging in psychology and neuroscience. Rather than attempting to resolve fast neuronal events directly, prior work has structured decision-making tasks into temporally separable stages. In particular, prolonging the decision phase allows rapid sensory responses to be dissociated from slower, decision-related ramping signals^33,34,44^.

Following this rationale, we designed the fMRI paradigm to exploit differences in the temporal profiles of sensory and decision-related signals. Our working hypothesis associated sensory responses with rapid BOLD response, and decision-related activities with slower accumulation of signal, producing a ramping hemodynamic signal. Accordingly, the sensory evidence presentation period was set to 6 s, matching the typical 4–6 s peak latency of visual-evoked responses in early visual cortex^45^, thereby improving SNR.

Immediately following the fixation period, a 2-second response epoch began, during which monkeys reported their decision via a saccade. The 6-second offset between the onsets of the evidence and response epochs is designed to temporally separate their respective BOLD signatures at mesoscale. A 10-second inter-trial interval followed to allow the BOLD signal to return to baseline. This temporal structure allows us to leverage 7T spatial precision to dissociate how distinct mesoscale modules contribute to sequential stages of the decision process.

This work represents a pioneering application of mesoscale fMRI mapping in awake, behaving macaques at 7T. Achieving submillimeter functional resolution under these conditions required overcoming substantial challenges, including physiological motion, fixation stability, and the design constraints of surface coils. We addressed these by combining extensive behavioral training and a custom sphinx-position chair with a secure headpost holder. Functional data were acquired without spatial smoothing and analyzed using T-statistics corrected for multiple comparisons with the false discovery rate (FDR) procedure, allowing reliable detection of fine-scale brainwide activation patterns. These procedures for revealing mesoscale functional domains (e.g. V2 color and motion stripes, V4 surface and contour domains, MT motion direction domains) in ventral and dorsal visual pathways using 7T fMRI were initially developed in anesthetized monkeys^16^ (refs), adapted for awake fixating monkeys (refs), and then further adapted for monkeys performing 2AFC tasks.

As shown in Fig. 1d, patchy activations were evident across early visual cortex (V1, V2, V4, and MT), and were consistent with retinotopy^17^. We then adapted these methods during performance of the visual discrimination task. Using an 8-channel customized RF coil (see Methods), BOLD responses were obtained in 7T fMRI during behavioral performance of the contrast discrimination task. The mesoscale BOLD timecourses from these activations in visual cortex demonstrate an expected contrast dependence of the BOLD response (Fig. 1e), reflecting a basic aspect of sensory encoding in visual cortex. As shown in Fig. 1f, the averaged peak responses (obtained from 6-8 sec period) exhibit canonical contrast response functions well-fit by a Naka-Rushton model^47^. This indicates that the mesoscale fMRI mapping during the contrast discrimination behavior is sufficiently sensitive to provide a quantitative basis for contrast encoding. We next examined whether these signals may also reflect decision making and action aspects of this task.

### Functional Networks Identified from Temporal Profiles

Our aim is to investigate spatiotemporal dynamics of task-engaged brain-wide networks. Given the temporally segregated task structure with distinct stimulus presentation (2-8 sec) and response (8-10 sec) phases, building on established methods for dissociating neural processes via distinguishing temporal profiles of hemodynamic response^24,25,39^, we modeled brain-wide task-phase-specific activity using separate regressors with a general linear model (GLM).

The fixation regressor was aligned to stimulus onset and captured activity during the 2–8 s interval (Fig. S2). The response regressor was time-locked to eye-movement onset and captured saccade-related activity during the 8–10 s interval. Although the 2–8 s period is referred to here as the fixation phase, it may encompass multiple processes occurring before the eye movement, including sensory processing, evidence accumulation, decision formation and motor preparation.

Specifically, we identified voxels significantly modulated by the fixation regressor (*FDR-corrected p < 0.01*). For each significant voxel, the time course was averaged across all trials and conditions, and linear regression was then used to estimate its temporal slope during the 2–8 s interval. The resulting slopes varied widely across voxels, reflecting differences in their temporal response profiles. Unsupervised clustering of these voxel-wise slope values using a Gaussian mixture model (GMM) revealed a robust bimodal distribution. Voxels were therefore divided into steep- and shallow-ramping groups using monkey-specific slope thresholds (monkey M: 0.100; monkey J: 0.065), corresponding to groups A and B, respectively. Group C comprised voxels significantly modulated by the response regressor (*FDR-corrected p < 10^⁻^*^6^), and showed a distinct population of movement-locked responses.

Following this characterization, all significant voxels were classified into three distinct groups based on their temporal dynamics. In monkey M, 455, 1,782, and 3,617 voxels were assigned to groups A, B, and C, respectively; the corresponding numbers in monkey J were 3,107, 1,646, and 4,920 voxels. The three groups exhibited distinct temporal profiles, as illustrated in monkey M (Fig. 2). Group A showed a rapid increase in BOLD signal following stimulus onset, consistent with sensory-related activity (Fig. 2a). Group B showed a more gradual increase during the fixation period, consistent with decision-related activity (Fig. 2b). By contrast, group C showed a delayed rise following response onset (8 s; dashed line), consistent with action-related activity (Fig. 2c). Each trace represents the mean time course across voxels within the corresponding group. Together, these distinct temporal profiles dissociated three functionally defined voxel populations associated with sensory processing, decision formation and action execution, providing a temporal basis for mapping their spatial organization across the brain.

### Spatial Transition of the Three Brain-wide Networks

Having identified three functionally distinct response groups, we next investigated how these temporally defined groups (A, B, C) are spatially distributed across the cortical surface, allowing us to trace the brain-wide transformation from sensation to action. The global map (Fig. 2d-f) reveals a large-scale, topographic shift: group A (Fig. 2d, red) responses were confined to early visual cortex, group B (Fig. 2e, yellow) to associative regions such as IPS and PFC, and group C (Fig. 2f, blue) to premotor and PFC territories. In visual cortex, group C also exhibited a shift in visual topography. Then, we overlay the three networks to detail this transition across functional systems (Fig. 2g-i).

The cortical processing begins in early visual cortex. During the fixation period (corresponding to group A, Fig. 2d), activations were seen in the retinotopically corresponding area at approximately 4° eccentricity due to the 2° diameter stimulus centered at 4.2° eccentricity (red patches labeled as V1, V2, and V4). Another laterally located red patch observed near the foveal representation (red patch labeled as Fovea) consistent with responses to the central fixation cue (0.2° radius).

Group B also participated in the fixation period and exhibited gradual ramping activity during the fixation phase, although with a more fragmented spatial organization (Fig. 2e). Group B voxels were distributed in the intraparietal sulcus (area LIP, yellow patch in Fig. 2g) and the superior temporal sulcus (area MT, FST). Furthermore, group B voxels were concentrated in the lateral prefrontal cortex, in ventral area 44 and 45b (yellow patches in Fig. 2h); they were also localized anteriorly, near the arcuate sulcus, within dorsal premotor areas F4 and F5 (yellow patches in Fig. 2i).

Group C, associated with the response period, exhibited increased activity following saccade onset. While group C voxels were also primarily distributed within regions such as IPS, premotor cortex, and PFC (Fig. 2h), they exhibited a topographic reorganization relative to group B. As shown in Fig. 2g, in the intraparietal sulcus, group B voxels are located more anteriorly than group C (yellow arrow indicates the direction from anterior to posterior).

In PFC, group C voxels extended dorsally across 45a/b, 8Av, and into ventrolateral regions including 46v and 46f (Fig. 2h). The resulting distribution was distinct from that in either group A or group B, and formed a mosaic-like layout spanning the arcuate and principal sulci.

In premotor and motor cortex, functional groups B and C (yellow and blue) also exhibited distinct anterior–posterior distributions (Fig. 2i). group C voxels, which are aligned to saccade onset, were situated more at the posterior bank of the central sulcus (3a and F1). We observed activation in area 3a, the proprioceptive representation of eye position in monkey primary somatosensory cortex^48,49^.

Overall, these data provide a multi-areal view of spatial activations over the course of the discrimination task. We note that the voxels in groups A, B, and C largely exhibited little overlap. In some areas, the two response types did show partial spatial overlap. For example, there was some overlap between yellow and blue in area F4, forming an intermediate zone between premotor and primary motor territories. However, overall, the overlap was minimal (Dice coefficient: group A–C = 0.02; group B–C = 0.14), forming interdigitated patches of spatially distinct group A, group B and group C activity, potentially relating to sensory, decision and action, respectively. The resulting overall network activations displayed shifts from (1) predominantly visual areas (group A, V1-V4) to (2) parietal (IPS areas such as LIP, VIP), prefrontal (44, 45b), and premotor (F4, F5) areas to (3) motor (F1) and prefrontal (FEF, 46v) areas.

Finally, we summarized the brain-wide distribution of these three groups in a region-by-category matrix based on the D99 macaque atlas (Fig. 2j). Each column represents one anatomically defined brain area, and each row reflects its assigned group. Areas were color-coded by system type and sorted according to their group affiliation.

For visualization, we report results from the hemisphere contralateral to the visual field containing the test stimulus and include both cortical and subcortical structures. Fixation-related voxels were further assigned to two cluster-defined groups. Using monkey M as an example, group A comprised 52 cortical areas and 31 subcortical areas; group B comprised 85 cortical areas and 54 subcortical areas. Response-related voxels constituted group C and comprised 99 cortical areas and 61 subcortical areas. Anatomical areas were defined according to the D99 atlas, which comprises 149 cortical and 219 subcortical areas.

### Distributed Representation of Choice Across a Mesoscale Network

To determine how the BOLD signals in mesoscale activation patches encode the animal’s upcoming behavior and whether they exhibit choice-related dynamics analogous to single-neuron recordings, we first tested a classical hypothesis: If an area carries a representation of choice (chosen versus unchosen for upcoming action), voxels within that area should exhibit stronger pre-response activity when the corresponding stimulus (in the contralateral visual field) is chosen than when it is unchosen. High-certainty trials allowed us to isolate this computation with minimal behavioral ambiguity. Consistent with this hypothesis, schematic responses (Fig. 3a, b) demonstrated robust chosen–unchosen separation (statistically difference in response magnitude), with elevated activity in the hemisphere contralateral to the selected target.

We first examined the sensory network (group A), which showed rapid, stimulus-locked responses during the fixation period (2–8 s). In early visual areas contralateral to the test stimulus (e.g., V1; Fig. 3c-f), BOLD amplitude scaled with contrast, consistent with classical contrast-response functions. Crucially, however, sensory representations of the fixed standard stimulus (50% contrast) also displayed reliable choice-dependent modulation. Despite the local physical input within the receptive field of standard stimulus remaining identical, BOLD signals in the hemisphere contralateral to the standard were significantly elevated when this stimulus was ultimately selected versus unselected (Fig. 3f). These results demonstrate that pre-response activity in the sensory cortex is dynamically modulated by the outcome of the bilateral stimulus competition.

We next asked whether the decision group (group B) exhibited signals indicative of a choice. Decision group including the IPS, exhibited chosen–unchosen separation during the fixation period (Fig. 3g–j), mirroring the ramping accumulation processes reported in electrophysiology. In both test- and standard-contralateral hemispheres, activity was reliably higher when the corresponding stimulus was ultimately selected, consistent with a representation that predicts the forthcoming choice.

We next examined whether the choice representation is maintained within the action-related neuronal response during the response period. The action group (group C) displayed a distinct and saccade-aligned response profile. In the ventral central sulcus, we see corresponding action activity topographically reflected in the hemisphere contralateral to the chosen target rose sharply after eye movement (Fig. 3m, n). These dynamics complete a sequential progression across the three networks: early sensory encoding, decision-predictive integration, and finally action implementation, revealing a full cortical trajectory from sensory evaluation to motor execution.

To quantify choice representation across the entire brain, we performed a standardized comparison of chosen versus unchosen trials for all atlas-defined areas. Areas were included in the analysis only if they met the activation criterion of at least three significant voxels, with *FDR-corrected p* < 0.05 for sensory and decision groups and *p* < 0.01 for the action group. Statistical comparisons were performed via two-sample Welch’s t-test (*p* < 0.05). For cross-area visualization, effect sizes were quantified using Hedges’ g, providing a standardized measurement of choice-related activity.

This analysis was conducted for the period prior to the response, spanning both sensory and decision groups. This summarized Hedges’ g matrix (Fig. 3o) reveals that predictive choice representation is not confined to local regions but is instead distributed across a brain-wide network. During the pre-response epoch, robust choice representation signals appeared not only in visual and parietal cortices but also in prefrontal, premotor, and subcortical structures. While most previous electrophysiology studies have focused on one or a few areas, this widespread pattern indicates a highly distributed choice representation instantiated across a broad mesoscale network. Similar patterns were observed in a second monkey (monkey J, Fig. S3), with consistent chosen–unchosen lateralization, confirming that the distributed architecture is robust across individuals.

### Mesoscale certainty-dependent cortical maps reveal choice encoding under uncertainty

Having established that voxel masks derived from high-certainty conditions reliably separated chosen from unchosen responses when sensory evidence was strong, we next asked what the activation profiles are for uncertain conditions. Specifically, we asked whether different levels of certainty could be encoded by the same spatial mesoscale network but graded in response amplitude, or by mesoscale networks residing within the same cortical areas but occupying separate spatially distinct mesoscale domains. The latter would be consistent with representation with sensory cortical areas, in which different parameters are represented in systematic maps and values encoded by spatially distinct locations within those maps. Distinguishing between these two models—amplitude modulation within a single network versus spatially distinct mesoscale networks—is essential for understanding both temporal and spatial aspects of decision-making architecture.

To examine how perceptual certainty is organized across the cortex, we first constructed a winner-take-all (WTA) map, in which each voxel is assigned to the contrast condition that elicits its strongest GLM activations. Specifically, for each contrast condition, we computed a separate GLM, thresholded voxel-wise *FDR-corrected p* < 0.05, and labeled each voxel according to the condition producing its minimum q value. Projecting these voxel labels onto the cortical sheet revealed interleaved clusters of WTA maps spanning brain regions (Fig. 4a). According to behavior, large contrast differences (e.g., 90% vs. 50% contrast) produced highly certain choices (>90% performance), whereas near-threshold contrasts (47–53% contrast) approached chance (∼50% performance). Grouping contrast levels into “uncertain” (47%, 50%, 53% contrast) and “certain to test side” (90% contrast) regimes therefore allowed us to visualize certainty-dependent mesoscale structure across cortex: voxels maximally driven by uncertain conditions were colored green, and those driven by certain conditions were colored red (see Fig S4 for full map). For this analysis, we compared the voxels responding best to certain chosen condition 90% (Fig. 4b, dark-red, ‘certain’ voxels) with those responding best to the uncertain conditions 47% and 53% (Fig. 4b, green, ‘uncertain’ voxels).

We next evaluated whether voxel groups identified as certain (red) or uncertain (green) could predict behavioral choices from their BOLD responses during certain trials (test stimulus=90%) and uncertain trials (test stimulus=47%/53%) in Fig. 4a. We designated 47% and 53% contrasts as uncertain trials, because these two uncertain contrasts jointly yielded trial counts comparable to the certain conditions. The 50% contrast condition was excluded: including it would introduce a third uncertainty level, resulting in an unbalanced design relative to the certain conditions and therefore biasing statistical comparisons.

We found that in several regions uncertain voxels selectively encoded choices during uncertain conditions, whereas certain voxels encoded choices during certain conditions, revealing a spatial dissociation of choice encoding across levels of certainty. For example, in the intraparietal sulcus (IPS), a key decision-related region, voxels dominated by the 90% contrast condition (certain, dark red) reliably predicted choice during certain trials, as shown in Fig4c: their BOLD amplitudes were significantly higher for trials in which the test stimulus was chosen than unchosen when sensory evidence strongly favored the test stimulus (90% chosen test vs. 10% unchosen test). *However, these same voxels carried no significant chosen–unchosen difference during uncertain trials (47% and 53%). Thus, ‘certain’ voxels carry choice-selective signals primarily when sensory evidence is strong.* In contrast, voxels dominated by 47% and 53% (uncertain voxels, green) displayed the opposite pattern (as shown in Fig4d). During uncertain trials, these voxels exhibited a robust separation between chosen and unchosen responses, yet during certain trials their responses did not significantly differentiate. Additionally, the strength of response in certain trials and uncertain trials were comparable (Fig S5). *In other words, uncertain voxels selectively encode choice information when sensory evidence is near threshold*.

This dissociation reveals that certainty and uncertainty are represented by distinct mesoscale subdomains within IPS: certain voxels support choice encoding under strong evidence, whereas uncertain voxels remain informative even when the stimulus is ambiguous. The matrix in Fig. 4e summarizes all regions exhibiting this pattern, encompassing much of the dorsal sensorimotor pathway, including MT, V3, parietal areas 5 and 7, LIP, premotor area F5, prefrontal area 46d, and orbitofrontal area 12o, and extending into the cerebellum, notably cerebellar lobules IV and V. Together, these findings show that *uncertainty engages an additional spatially distinct mesoscale network that remains predictive even when sensory evidence is weak*.

## Discussion

### From fragmented findings to a whole-brain map of perceptual decisions

Perceptual decision making has been extensively studied using electrophysiology and behavioral modeling, yielding detailed insights into temporal dynamics such as evidence accumulation and decision commitment. Although recent studies have established the distributed nature of decision-related activity^7^, how this activity is spatially organized within and across brain regions remains incompletely understood. Here, by combining a perceptual decision paradigm with ultrahigh-field (7T) fMRI in awake macaques, we move from studies focused on a small number of regions toward a whole-brain map of perceptual decision making within individual subjects. Our results reveal task-related activations organized into mesoscale domains across the brain, providing a detailed map of distributed decision-related networks. Thus, while increasingly, studies of perceptual decision making in primates are conducting simultaneous recordings from multiple areas^3,40–44^, such as the middle temporal area (MT) and lateral intraparietal area (LIP), our study uses fMRI to reveal the brain-wide mesoscale network. We recognize that during this task, there may be multiple behavioral components (e.g. reward related, prior information) that contribute to these activations. However, mapping at brainwide scale does provide a distinct view of the brain state, in which shifts in multi-areal activity underlie task-related behavioral phases of sensory processing, decision formation and action.

Beyond this, our data suggest that decision-related information is also expressed through the spatial organization of mesoscale domains within cortical areas. Whereas previous studies have examined subspaces or manifolds of neural activity in locally sampled populations^5^ and changes in population activity across levels of sensory evidence^45^, our findings highlight that distinct mesoscale spatial domains within cortical areas carry decision-related information (e.g. degree of certainty^46^) under different task conditions. Preserving mesoscale spatial resolution allows condition-dependent response differences to be revealed using general linear models and simple conditional contrasts, without requiring complex modeling. These findings suggest that mesoscale spatial codes provide an additional dimension for understanding how distributed activity supports perceptual decisions.

### How do slow BOLD signals reflect to rapid decision making events?

Hemodynamic BOLD signals are events in the brain that extend over 10 sec, and so do not track the rapidly occuring neural events during a task that typically spans only hundreds of milliseconds. While it is the onset of specific sensory and behavioral events that trigger BOLD onset, this information is not enough. However, when these BOLD signals are further evaluated for spatial location, the BOLD specificity then emerges. That is, single mesoscale loci exhibit task-phase specific BOLD response, leading to the observed distinctions between three groups of timecourses associated with sensory, ‘decision making’, and response phase. In our task, the during the 6 sec fixation phase, both sensory and ‘decision making’ events are occurring; as shown in Fig.,.., these events can be distinguished by fast rise time/high amplitude profiles and slow rise time/low amplitude BOLD profiles, reflecting early sensory events followed by ongoing ‘decision’ events. Moreover, when these fast and slow groups are mapped separately, they associate largely with mesoscale domains in sensory (V1-V4) and higher order parietal (LIP) and frontal areas known to be associated with sensory and decision making phases. The latest response phase is clearly temporally separable and associates with eye movement related areas such as FEF^47^. Thus, although the BOLD signal is slow, it can be useful by isolating the combined spatiotemporal event that distinguishes the activity of different sets of mesoscale domains.

### Where and how are decision variables organized at mesoscale?

A central open question in cognitive neuroscience concerns the spatial encoding of decision variables across the brain. While classical models emphasize temporal computations—such as the accumulation of evidence—it remains largely unknown how these computations are embedded in the spatial architecture of the brain. In particular, widespread decision-related activity raises the question of whether individual areas contain spatially distinct domains that represent values of specific behavioral axes (such as certainty^48,49^).

By mapping task-related BOLD responses across the brain, we identify networks with distinct temporal profiles and mesoscale spatial organization. *Across all three functional networks, mesoscale BOLD signals consistently anticipated the animal’s eventual choice.* The sensory-related group exhibited contrast dependence and choice modulation, even for physically identical stimuli (Fig3c-f). The decision-related group showed a gradual BOLD buildup during fixation, consistent with sustained decision-related processing, although the temporal resolution of BOLD precludes identifying neuronal evidence accumulation directly. The action-related group exhibited later, not surprisingly, lateralized responses associated with the saccadic report^22,50^. When extended across the entire brain, these signatures revealed a distributed mesoscale circuit in which choice encoding was expressed throughout visual, parietal, prefrontal, premotor, and subcortical structures (see matrix in Fig2). The action-related group exhibited lateralized responses associated with the saccadic report. Together, these observations indicate that decision variables are instantiated through coordinated activity across spatially distributed mesoscale domains.

### Local mesoscale network transitions underlie perceptual decision making

Area-based descriptions of perceptual decision making emphasize a broad progression from visual processing through parietal and frontal circuits to motor output. Alongside these large-scale shifts in network, we observed additional local shifts that has not previously been described. Specifically, we observed network transitions of mesoscale functional domains within each of the parietal, frontal, and motor cortical regions (Fig. 2g-i). These spatial differences corresponded to successive task phases, from stimulus viewing to the saccadic report. That is, within regions (such as intraparietal sulcus, prefrontal cortex, and premotor cortex), sensory-related, decision-related, and action-related signals occupied adjacent yet largely non-overlapping spatial domains.

Thus, a single anatomical region can contain multiple domains with different task-related response profiles, distinctions that may be obscured by averaging activity across the area and which may not be evident from single site electrical recordings. These findings demonstrate that perceptual decision making involves not only transitions globally, but also reconfigurations of mesoscale functional domains within parietal, frontal, and motor associated regions, revealing a finer spatial organization than is incorporated by area-based models.

### Decision variables are encoded in a distributed brain-wide network in primates

A further question is whether decision variables are confined to specialized regions or distributed across the primate brain. Large-scale recordings in rodents report widespread decision-related signals^25,27^; as well, in primates multi-areal recordings^24,31^ find distributed decision-related activity, suggesting a basis in brain-wide networks. Here, we provide a more systematically sampled look at such networks at brain-wide scale.

Our results provide direct evidence that decision variables are encoded in a distributed mesoscale network spanning cortical and subcortical regions, and, moreover, in both hemispheres. We found that choice-related BOLD signals extended across sensory, associative, and motor systems, expressed in a lateralized and task-phase–specific manner. Importantly, this distribution was highly specific: different areas contributed to different phases of the decision process and, even within single areas, distinct mesoscale domains contributed to encoding of different phases. Together, these findings indicate that perceptual decisions are not confined to canonical decision areas, but rather represented at spatially specific (mesoscale) and coordinated brain-wide networks.

### Perceptual certainty and uncertainty are encoded in distinct spatial patterns

Beyond the spatial extent of decision networks, our data address how these networks adapt to varying levels of perceptual certainty. Classical models typically associate uncertainty with attenuation of the amplitude of decision-related neural activity. Our findings, however, reveal a different mechanism. We show that certainty and uncertainty are encoded in distinct spatial patterns at mesoscale. Voxel populations that robustly encode choice under high-certainty conditions are spatially distinct from populations that encode choice at near-threshold (uncertain) conditions (Fig.4). Notably, uncertainty engaged a distinct mesoscale network reconfiguration that remained informative (distinguished chosen and unchosen) when sensory evidence was ambiguous. This result indicates that the previously described attenuation of activity with uncertainty does occur in domains encoding certainty. Thus, as a decision is being made, the brain-wide network shifts locally within an area (such as IPS) from a ‘certain’ mesoscale network to an ‘uncertain’ network. Together, these findings support a spatial component of decision-related coding that complements neuronal response amplitude. We are not aware of electrophysiological data that report increase in response for ‘uncertain’ conditions, the lack of which may be due to sampling issues^41,45^.

### Take-home message: the brain thinks in space as well as in time

In summary, our findings demonstrate that perceptual decisions emerge from distributed, brain-wide mesoscale dynamics. Decision variables are neither localized to single areas nor simply broadcast across multiple areas in the brain. Instead, they are instantiated through spatially organized modular networks that shift with task demands and perceptual certainty.

This organization raises a computational hypothesis: spatially selective pooling and differencing may allow task-relevant information to be read out through relatively simple operations. Selective access to different domains could support flexible behavior as sensory conditions change. Although simple statistical analyses do not establish the computations performed by the brain, these findings motivate tests of whether spatial organization enables computationally and energetically economical readout.

For brain–computer interfaces, this perspective motivates recording and decoding strategies that preserve mesoscale spatial information and accommodate condition-dependent changes in the location of informative signals. Such strategies could help maintain useful readouts across changing task conditions and provide a basis for investigating long-term decoding stability. Establishing this benefit will require longitudinal recordings and independent tests of decoding performance.

In this framework, decision-related computations unfold not only over time but also through structured spatial organization at mesoscale. These brain-wide mesoscale networks have been observed consistently in vision neuroscience and now link known underlying brain-wide networks to phases of behavior. These functional domains constitute the substrate through which sensory information is transformed into decisions and actions, establishing a fundamental organizing principle for perceptual decision making.

## Method

### Subjects

All experimental procedures were conducted in accordance with the National Institutes of Health Guide for the Care and Use of Laboratory Animals and approved by the Institutional Animal Care and Use Committee of Zhejiang University. Two adult rhesus macaques (Macaca mulatta) participated in this study: Subject M (male, 7-year-old, 6-8 kg) and Subject J (female, 11-year-old, 4-6 kg). Both animals were implanted with MRI-compatible headposts and trained to sit in a sphinx position in a custom-designed primate chair. Behavioral motivation was achieved through a regulated hydration protocol, with fluid rewards administered during experimental sessions and free water access provided weekly. Daily minimum hydration levels were systematically maintained while monitoring health parameters including body weight, food intake, and excretory patterns.

### Behavioral training and task paradigm

Monkeys were trained over several months to perform a two-alternative forced choice (2AFC) contrast discrimination task. Training included three stages: (1) motion control via body-still training using a 3D accelerometer-based feedback system, (2) fixation training using a mock scanner with real-time eye tracking (Eyelink 1000 plus, CR system, Canada), and (3) acclimatization to the 7T scanner environment. Initial scan training was aided with a mild sedative (dexmedetomidine, gradually reduced from 20μg/kg to 8μg/kg).

Each trial commenced with a 2-s central fixation period, followed by a 6-s bilateral presentation of grating stimuli. The sinusoidal gratings were rendered as circular patches (2° diameter) positioned at matched eccentricities (4.2°) in bilateral visual fields. The standard stimulus with fixed 50% contrast was presented in one hemifield, while the contralateral test stimulus contrasts varied pseudo randomly across trials (10-90% range). In a single day, the standard stimulus was maintained on one side and left/right counterbalanced across days. After 6-s stimulus presentation, the central fixation point disappeared, and the subjects were required to saccade to the higher contrast stimulus within 2-s and maintain a 0.5 s gaze to a 1 deg radius response window overlying the grating. Only correct responses were rewarded with juice. When the test stimulus was the same as the standard stimulus, both are considered correct answers. Trials were separated by a 10-s inter-trial interval (ITI). Eye position was monitored with an MRI-compatible infrared tracking system (Eyelink 1000 Plus; SR Research), sampled at 250 Hz.

Any fixation break (deviation beyond a 1° radius window) lasting longer than 1 s during the stimulus phase, or longer than 0.5 s during all other task phases, triggered immediate trial abortion. Aborted trials were followed by the presentation of a uniform gray screen until the initiation of the subsequent trial.

Only trials with stable fixation during stimulus presentation (with no fixation break exceeding 1 s within a 1° radius window) and valid behavioral responses (saccadic choice was made during the response window) were included in analyses. A total of 18 fMRI sessions were conducted across both monkeys, yielding 1,761 valid trials (monkey M: 875 trials; monkey J: 886 trials). One example dataset from monkey M (5 sessions, 466 valid trials, left side as the standard stimulus configuration) is shown in Fig. 2b.

### Visual stimuli

Visual stimuli were generated using Psychopy^50^ and presented on a translucent screen positioned 60 cm in front of the animal via an MRI-compatible projector (PROPixx, VPixx Technologies, Canada; resolution: 1920 × 1080 pixels; refresh rate: 60 Hz RGB, 1440 Hz grayscale). The visual stimuli consisted of two sinusoidal gratings patterns presented as circular patches (2° diameter) symmetrically positioned in the lower visual fields at 4.2° eccentricity along 225° (left) and 315° (right) meridians. Gratings maintained vertical orientation with spatial frequency fixed at 1 cycles/degree and temporal frequency fixed at 3Hz.

Visual stimuli were presented on a gamma-corrected display, such that digital input values were linearly mapped to physical luminance. Screen luminance was calibrated using a photometer, and the mean background luminance was maintained at 200 cd/m^2^ throughout the experiment. Stimulus contrast was quantified using Michelson contrast, defined as

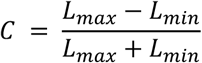

Where *L_max_* and *L_min_* denote the maximum and minimum luminance values of the sinusoidal grating, respectively.

The contrast of the test stimuli varied across seven conditions: 10%, 40%, 47%, 50%, 53%, 60%, and 90%. All contrast conditions were pseudorandomly interleaved to minimize adaptation effects.

### MRI acquisition

Scanning was performed on a 7T whole-body MRI system (Magnetom, Siemens Healthcare), with custom-built RF coils optimized for mesoscale functional imaging. An 8-channel customized whole-brain coil (1 mm isotropic) was used for functional scans during the contrast discrimination task.

To improve tSNR through enhanced SNR homogeneity, facilitate parallel imaging, and accommodate multimodal MRI/fMRI study devices, the coil design features an 18-cm diameter transmit-only loop positioned approximately 1 cm above an array of eight receive-only loops (5–8 cm in diameter). The transmitter and receiver are effectively decoupled via active detuning circuitry, while the receive loops are decoupled through a combination of overlapping geometry and preamplifier decoupling. To provide sufficient space for headpost fixation and multimodal device integration, each preamplifier is situated approximately 51 cm from its respective receive element. Furthermore, lattice baluns at each element’s drive point eliminate common-mode currents. The coil was fabricated on a 3D-printed polycarbonate (PC) former using Fused Deposition Modeling (FDM). Four 0.25-inch Loc-Line modular hoses allow for adjustable positioning. Finally, seven housing openings provide direct access to the occipital, temporal, and frontal lobes and accommodate the headpost.

Functional images were acquired using a T2*-weighted gradient-echo EPI sequence with the following parameters: TR = 2000 ms, TE = 25 ms, flip angle = 90°, resolution = 1 mm isotropic bandwidth = 1158 Hz/pixel. Each trial was treated as a single BOLD block. Anatomical images were acquired using a T1-weighted MPRAGE sequence (TR = 2500 ms, TE = 3.21 ms, resolution = 0.5 mm isotropic).

### MRI Data analysis

#### Preprocessing

Functional data were preprocessed using AFNI^51^ and FreeSurfer^52^. Steps included slice timing correction, motion correction, distortion correction (using reversed phase-encoded images), and co-registration to anatomical scans. Data were aligned to the sphinx position. *Individual-level statistical inference.* Statistical inference at individual level was implemented using a general linear model (GLM) framework via AFNI’s 3dDeconvolve. The GLM functioned as a multivariate regression to partition BOLD signal variance into task-related and confounding components. The observed time series (*Y*) was modeled as a linear combination of explanatory variables (*X*, the design matrix) and a residual error (*ε*): *Y = Xβ + ε*. The design matrix incorporated task-specific regressors (fixation, 2–8 s post-trial onset; response phases, 8–10 s post-trial onset) convolved with a canonical hemodynamic response function (HRF), supplemented by covariates of no interest (motion parameters and polynomial trends to account for low-frequency drift). To address the intrinsic temporal autocorrelation of fMRI noise, we further employed restricted maximum likelihood (REML) estimation via 3dREMLfit with an ARMA(1,1) model, yielding robust *β* coefficients and corresponding *t* statistics. To control for the inflated type I error rate associated with voxel-wise inference across the whole brain (∼10^5^ voxels), we applied false discovery rate (FDR) correction. FDR constrains the expected proportion of false discoveries among all rejected null hypotheses. For intuitive visualization, FDR-corrected *p* values were transformed into standard normal Z-scores based on the unit Gaussian distribution *N(0,1)*, satisfying the relationship *P(|x| > z) = p*. This non-linear mapping achieves range compression of the highly skewed *p* value distribution, ensuring that Z-scores are monotonically and positively correlated with statistical significance. It is important to distinguish this statistical Z-score—a probability mapping of significance levels—from the Z-score normalization applied during preprocessing, which scales the raw BOLD signal amplitude to unit variance.

#### Clustering of voxels

To further quantify fixation-period dynamics, trial-averaged timecourses for each voxel significantly modulated by the response regressor (*FDR-corrected p* < 0.01) were fit with a simple linear function, and the resulting slope was used as a summary measure of the voxel’s response profile. Voxels were clustered using a Gaussian Mixture Model (GMM) applied to their slope values, which identified two groups: a steep-slope cluster (group A, sensory-related) and a shallow-slope cluster (group B, decision-related). The Bayes-optimal intersection—defined as the slope at which the posterior probabilities of the two GMM-derived Gaussian components are equal—yielded subject-specific thresholds of 0.100 for monkey M and 0.065 for monkey J, separating steep from shallow fixation-period dynamics.

Voxels significantly modulated by the response regressor (*FDR-corrected p* < 10^-6^) were classified independently as group C (action-related). In this way, significant voxels were divided into three groups—sensory-related (group A), decision-related (group B), and action-related (group C)—as illustrated in Fig. 2 and Supplementary Fig. S1.

#### Statistical analysis of chosen versus unchosen responses

For each voxel, we quantified the extent to which BOLD activity predicted behavioral choice by comparing the BOLD response on trials where the monkey chose the stimulus in the contralateral visual field against trials where that stimulus on same location was unchosen. All analyses were performed on trial-wise timecourses extracted from voxels that showed significant modulation by the corresponding GLM regressors.

Analyses were performed at the voxel–trial level, where each data point corresponds to the full BOLD timecourse of a single voxel in a single trial.

For the remaining trials, BOLD responses were first averaged across significant voxels on a trial-by-trial basis. Mean responses were then computed within predefined temporal windows, including a broad stimulus window (2–8 s) and a decision-related window defined as either 6–8 s or 12–14 s, depending on whether the underlying GLM regressor was fixation-locked or response-locked.

Within each analysis window, we compared the BOLD response on trials where the monkey chose the stimulus in the contralateral visual field against trials where that stimulus on same location was unchosen, using two-sample Welch’s t-tests to accommodate unequal variances. Effect sizes were quantified using Hedges’ g, providing a bias-corrected standardized measure based on the pooled standard deviation. Ninety-five percent confidence intervals for g were estimated using non-parametric bootstrap resampling of trials (5,000 iterations with replacement). Unless otherwise stated, bar plots depict mean ± s.e.m., and statistical significance is indicated by asterisks (p < 0.05; *p < 0.01; **p < 0.001; ***p < 0.0001). To control for multiple comparisons across voxels, p-values were corrected using the Benjamini–Hochberg procedure, yielding false discovery rate corrected q-values.

These per-area effect sizes and statistics were used both for single-area predictive analyses and for constructing the area-by-system effect-size matrices shown in Fig. 3. In addition, a paired two-row matrix was generated to directly compare Hedges’ g values computed from the 6–8 s window for chosen and unchosen conditions, using an identical area ordering and a shared color scale.

### Behavior Data analysis

Psychometric curves were fit using a logistic regression, and contrast response functions were fit using the Naka-Rushton model. SEM across trials was computed for each contrast condition. All statistical analyses were performed using python.

## Supplementary Data

**Fig. S1.**
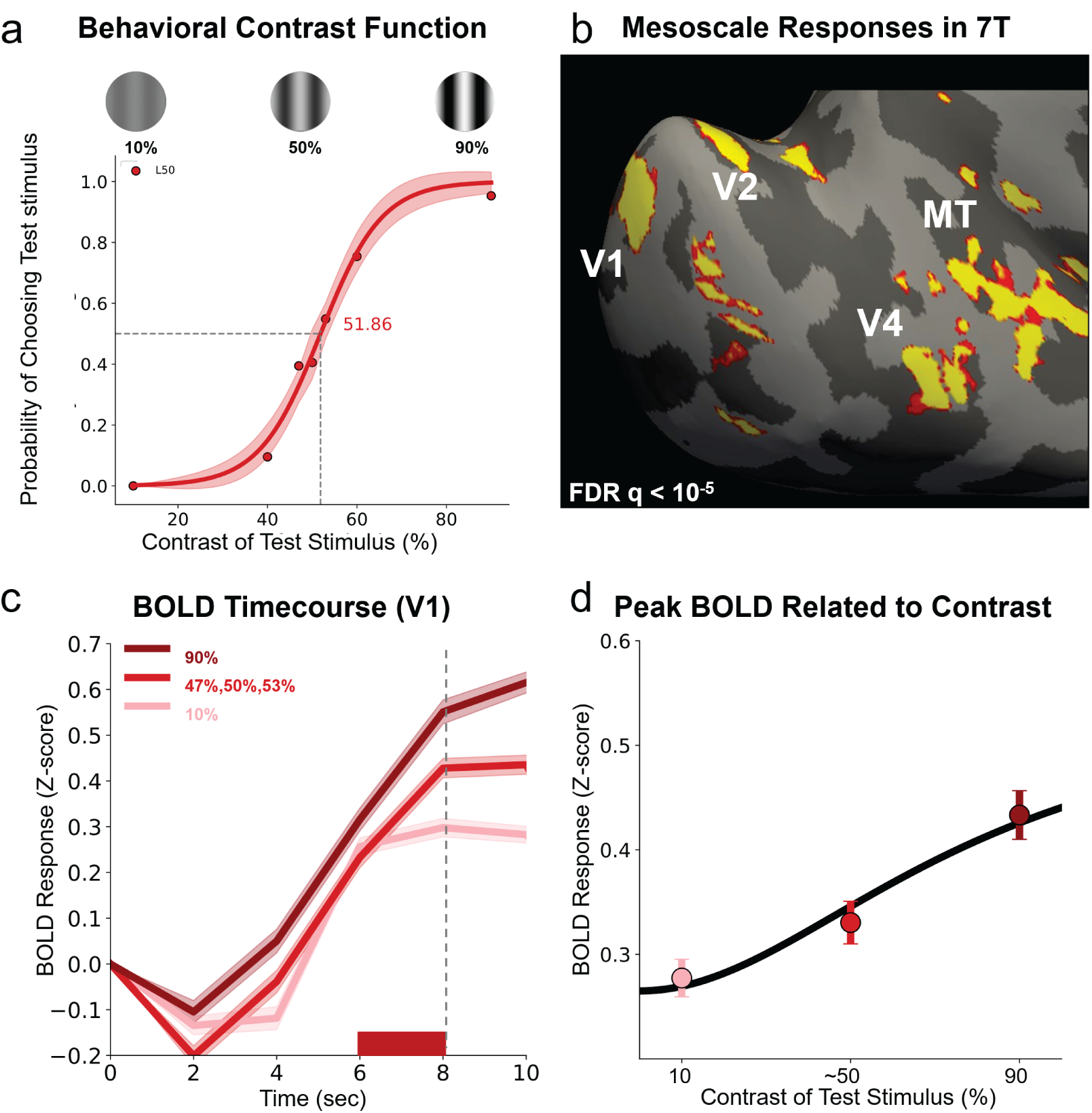
Behavior curve and BOLD response of monkey J. **a,** Representative psychometric function showing the probability of choosing the test stimulus (monkey J; 886 trials trials, the left side was fixed as the standard stimulus). The point of subjective equality (PSE) was estimated at 51.86% for monkey J, under conditions where the left side was fixed as the standard stimulus. Shadowed error bars represent the standard error of the mean (SEM) across all trials for each of the seven test contrast levels (10%-90%). **b,** Activation in mesoscale functional organization in early visual cortex mapped in 7T fMRI during a fixation task. (*FDR-corrected p* < 10⁻^5^) **c–d,** Contrast-dependent BOLD responses in V1 measured via mesoscale fMRI. **c**, Time course of BOLD signals from V1 across test contrast conditions. Responses increased monotonically with contrast, from pink (10%) through red (average of 47%, 50%, and 53%) to dark red (90%). Shaded areas denote SEM across trials. The gray dashed vertical line marks the onset of saccade initiation. **d**, Peak BOLD responses (averaged over 6–8 s, red horizontal bar in Fig. 3e) plotted as a function of test stimulus contrast. Data were well-fit by a Naka–Rushton function, capturing the characteristic compressive nonlinearity of cortical contrast gain. Error bars indicate the standard error of the mean (SEM) across trials. Valid trial counts at each contrast level were as follows: 65 trials at 10%, 67 at 90%, and 207 combined across 47%, 50%, and 53% contrasts. All data were from monkey J under the condition where the left side was fixed as the standard stimulus.

**Fig. S2.**
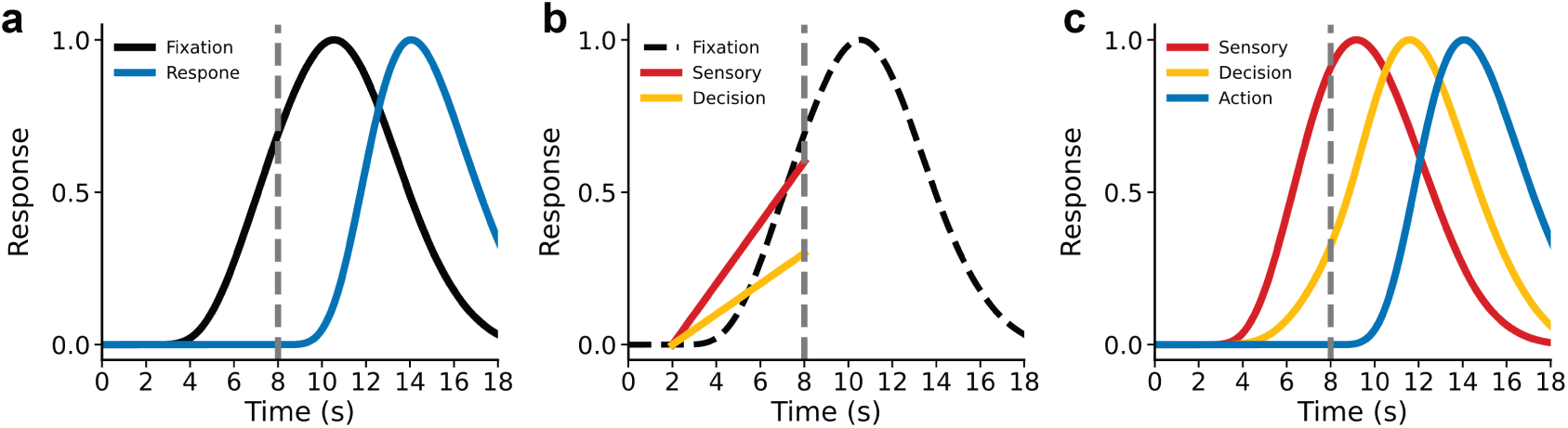
Time-resolved categorization of voxels reveals three functional networks. **a,** Illustration of the fixation regressor (black) and response regressor (blue) used in the general linear model. Fixation onset and response onset were each convolved with a canonical hemodynamic response function (HRF). Only significantly activated voxels were included in subsequent analyses. Statistical significance was assessed using false discovery rate correction (fixation regressor: *FDR-corrected p* < 0.01; response regressor: *FDR-corrected p* < 10^-6^). **b,** Voxels in the fixation group were further clustered based on the rate of BOLD signal increase, revealing two distinct temporal groups: a rapid-slope group (red, sensory-related, group A) and a slow-slope group (slow, decision-related, group B). **c,** Together with the action-related response timecourses (blue, group C), this procedure yielded three major functional groups of timecourses during perceptual decision-making.

**Fig. S3.**
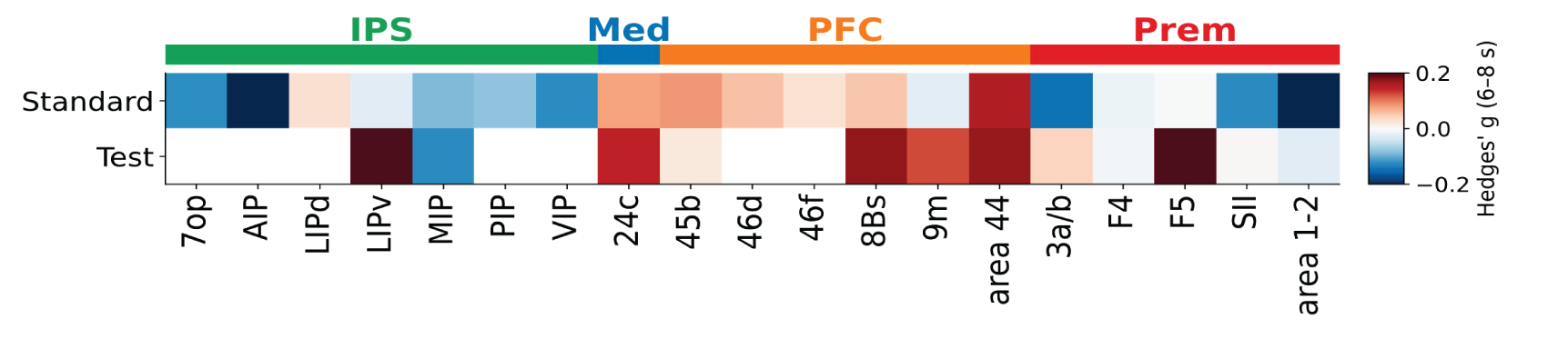
Reproducible spatial organization of choice representation across animals. Choice representation in an independent animal (monkey J). Summary matrices of chosen–unchosen effect sizes (Hedges’ g, 6–8 s) across all areas. Each column corresponds to one area defined by D99 atlas, grouped and color-coded by functional system. Color intensity reflects the magnitude of the chosen–unchosen difference, quantified using Hedges’ g computed within the 6–8 s post-stimulus window. System labels are abbreviated as follows: Med, medial; Prem, premotor; Subc, subcortical. *FDR corrected p* < 0.05. Despite differences in individual sessions and sampling statistics, a qualitatively similar choice representation matrix structure was observed in monkey J, indicating a consistent mesoscale organization of choice-related signals across animals.

**Fig. S4.**
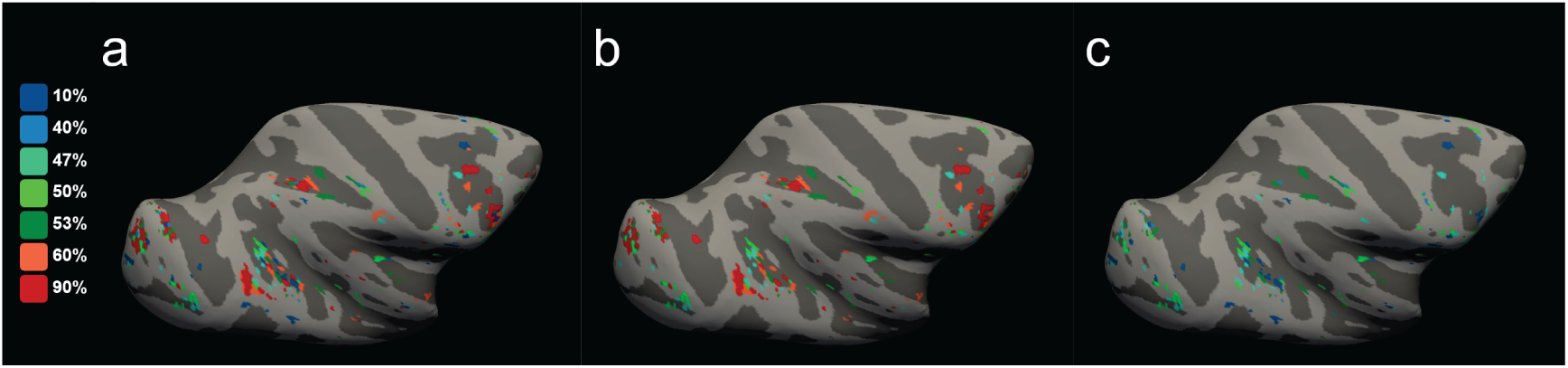
Winner-take-all cortical maps across all contrast conditions. Winner-take-all maps were generated by assigning each cortical vertex to the condition that elicited the maximal response amplitude. **a,** WTA map computed using all contrast conditions. **b,** WTA map computed using trials with uncertain contrasts (47%, 50%, and 53%) together with high-certainty conditions favoring the test side (60% and 90%). **c,** WTA map computed using trials with uncertain contrasts (47%, 50%, and 53%) together with low-certainty conditions favoring the test side, high-certainty conditions favoring the standard side (10% and 40%).

**Fig. S5.**
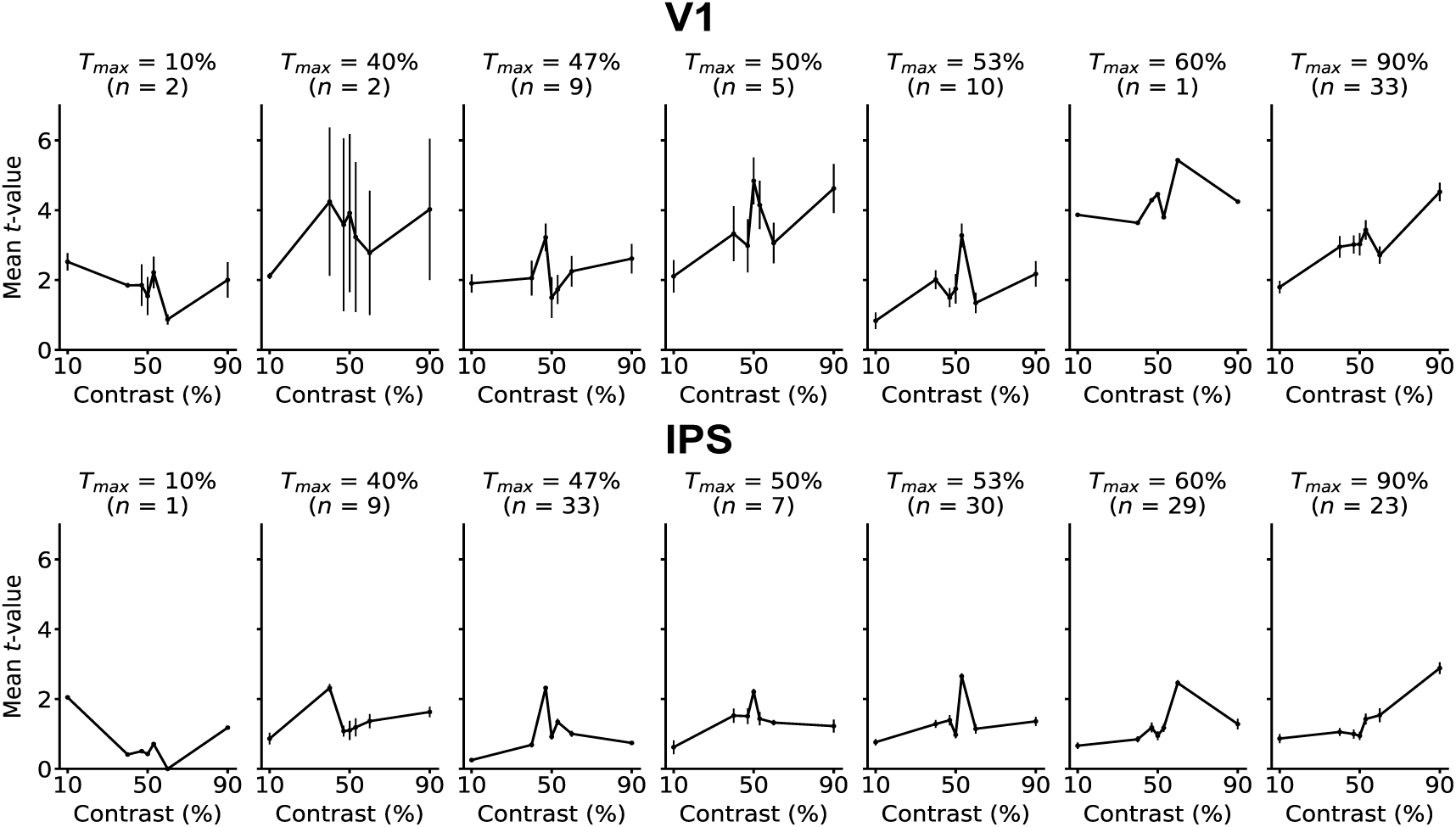
Uncertain preference distribution of V1 and IPS. For each of the seven contrast conditions (1–7, from lowest to highest), we identified voxels whose most significant activation (maximum T-value in voxel-wise FDR-corrected GLM) corresponded to that condition. These voxels are referred to as “T-max = condition n” groups. For each group, we extracted the average t-statistic across all voxels in the group for every contrast condition, generating a preference distribution. top row: uncertain preference distribution of V1; bottom row: uncertain preference distribution of IPS. Each column shows the curve for a different t-max group (condition 1 to 7, left to right). Error bars or shaded areas indicate s.e.m. across voxels.

